# Early menopause is associated with reduced global brain activity

**DOI:** 10.1101/2025.10.10.681622

**Authors:** Xufu Liu, Liying Luo, Laura Pritschet, Yutong Mao, Feng Han, David N. Proctor, Xiao Liu

**Affiliations:** Department of Biomedical Engineering, The Pennsylvania State University, University Park, PA, 16802, USA; Department of Sociology and Criminology, The Pennsylvania State University, University Park, Pennsylvania, 16801, USA; Department of Psychiatry, University of Pennsylvania, Philadelphia, PA, 19104, USA; Department of Kinesiology, The Pennsylvania State University, University Park, PA, 16801, USA; Intercollege Graduate Degree Program in Integrative and Biomedical Physiology, The Pennsylvania State University, University Park, PA, 16801, USA; Institute for Computational and Data Sciences, The Pennsylvania State University, University Park, PA, 16802, USA

**Keywords:** Early menopause, Global brain activity, Resting-state fMRI, Infra-slow brain activity

## Abstract

Menopause affects the aging process in women through significant ovarian hormone production decline in midlife. Women who experience early menopause face an accelerated physiological aging rate, along with impaired memory and increased risks of neurodegenerative diseases. However, it remains elusive how the timing of menopause affects brain activity, which could be crucial for understanding menopause-related acceleration of aging and increased risk of dementia. Recent studies have revealed a highly structured infra-slow (< 0.1 Hz) global brain activity across species and linked it to arousal and memory functions, as well as waste clearance in Alzheimer’s diseases (AD). In this study, we examined how this global brain activity relates to age of menopause using resting-state fMRI data from the Human Connectome Project-Aging dataset. We found that women who experienced earlier menopause (mean menopausal age 45±3.5 yr) exhibited weaker global brain activity (*p* = 5.0*×*10^-4^) with reduced coupling to cerebrospinal fluid (CSF) flow (*p* = 0.017) compared to age-matched later-menopausal women (mean menopausal age 54±1.2 yr). Differences appeared mainly in higher-order brain regions, where activation levels correlated with memory performance in earlier but not in intermediate or later menopausal women. These findings highlight brain activity changes linked to early menopause, suggesting a potential mechanism underlying memory decline and the increased risk of AD and dementias in early-onset menopausal women.

## Introduction

Menopause, clinically defined as twelve consecutive months without a menstrual period, marks the permanent cessation of ovarian function and is accompanied by a decline in estrogen and progesterone in the middle decades of life^1,2^. Given their role in regulating neurotransmission, synaptic plasticity, and neuroprotection^3–5^, the loss of these steroid hormones can trigger brain reorganization and speed up the aging process^6–9^. The age in which a woman experiences menopause is a critical factor in accelerating the brain aging ^4^. Premature (35-40) and early menopause (40-45) have been associated with an increased risk of cognitive and memory impairments, as well as dementia^10–13^, particularly Alzheimer’s disease (AD)^14^. The decline in estrogen following menopause, along with its subsequent effects on neuromodulation-particularly on the cholinergic system-has been hypothesized as a potential pathway underlying menopause-related brain changes^15–18^. However, it remains unclear how the timing of menopause affects brain activity, which could be crucial for understanding neural mechanisms underlying the brain aging in midlife, as well as increased susceptibility to neurodegeneration among women.

Recent research has revealed a highly-structured, infra-slow (<0.1 Hz) global brain activity, reflected as large peaks in the global mean fMRI blood-oxygenation-level dependent (gBOLD) signal, and linked this activity to arousal and memory functions^19–22^. In both human fMRI and monkey electrocorticography (ECoG) data, this activity manifests as global waves propagating across cortical hierarchies between lower-order sensory-motor (SM) areas and higher-order brain networks, particularly the default mode network (DMN)^23,24^. These global waves are also associated with specific de-activation of the Nucleus Basalis (NB) in the basal forebrain, a key hub of cholinergic neurons known to influence memory functions^25,26^. The infra-slow global brain activity was observed in large-scale neuronal recordings from mice as stereotypic cascades of sequential activations across two distinct groups of neurons^22^. This global dynamic is accompanied by changes in the rate of hippocampal sharp wave ripples (SPW-Rs) and arousal indicators, such as delta-band (1-4 Hz) activity or pupil size, suggesting its connection to the arousal and memory systems^22^. The spiking cascade is also synchronized with hippocampal replay events^21^. Intriguingly, the spiking cascades and fMRI waves are coordinated with opposite modulations in sensory encoding and memory process/retrieval across their cycles in mice/humans^20,21,27^.

The infra-slow global brain activity has also been linked to neurodegenerative diseases and aging^28–31^. The gBOLD activity is much stronger during light sleep and coupled with cerebrospinal fluid (CSF) flow^32^, suggesting its potential link to sleep-dependent, CSF-related perivascular clearance^33^. Consistent with this hypothesis, the strength of gBOLD-CSF coupling was subsequently found to correlate with various AD pathologies, particularly the accumulation of amyloid-beta (A*β*)^30^, as well as cognitive decline in Parkinson’s disease (PD)^29^. In addition, the propagation of gBOLD waves has been shown to explain the early spreading pattern of A*β*^28^. Changes in gBOLD-CSF coupling appear to extend beyond pathological conditions and during normative aging. It begins to decline around age 55, particularly in women^31^, as shown in healthy cohorts from the Human Connectome Project Aging (HCP-A)^34^. Interestingly, age and menopause status showed a notable interaction effect on the coupling measure, with postmenopausal women exhibiting a faster decline between age 35 and 55 compared to pre and perimenopausal women^31^. This finding highlights the potential relevance of menopause on gBOLD activity, including it’s potential contribution to the increased risks of dementia and memory impairments. What remains unclear, however, is whether the age at which menopause affects this gBOLD activity and its coupling with CSF flow decades after this transition state. If so, do these differences in gBOLD activity account for memory declines in women who experience menopause earlier than average?

To answer these questions, we used resting-state fMRI data from postmenopausal women in the HCP-A project to examine the impact of one’s age at menopause on infra-slow global brain activity, as measured by gBOLD signal, focusing on its spatiotemporal patterns, coupling with CSF flow, and connection to memory functions. We observed weaker gBOLD activity, as well as its weaker coupling with CSF, in women who experienced menopause relatively earlier (before age 49) compared to those who experienced it later (after age 52), with the most pronounced differences in higher-order brain areas, particularly the DMN. Importantly, weaker activations of gBOLD waves in these higher-order brain regions are associated with poorer performance in memory tasks among earlier menopausal women. These findings highlight brain activity changes related to early menopause and suggest their potential contribution to the increased risk of memory impairments and dementia observed in women with early menopause.

## Results

### Earlier menopause shows weaker gBOLD activity and gBOLD-CSF coupling

We focused on 90 female subjects from a total of 124 HCP-A subjects with menopause-related reproductive health data, excluding 34 individuals outside the age range of 52.9 to 66.5 years to minimize the confounding with age at MRI (see Methods and **Fig. S1** for more details). These 90 subjects were then divided into three equally sized groups based on their age at menopause (**Fig. 1A**): earlier (age: 38.0-49.2, 45.5±3.5 yrs), intermediate (age: 49.2-51.7, 50.4±0.6 yrs), and later (age: 51.7-56.4, 54.0±1.4 yrs) menopausal groups. The three groups exhibited no significant differences in age at MRI (earlier: 58.1± 3.7 yrs, intermediate: 58.2± 4.0 yrs, later: 60.0± 3.7 yrs; *p* = 0.11, ANOVA; **Fig. 1B**), years of schooling (*p* = 0.15), or body mass index (BMI) (*p = 0*.*18)* (**Table 1**). Compared to later menopausal women, earlier menopausal women exhibited significantly weaker (i.e., less negative) gBOLD-CSF coupling (*p* = 0.017, two-sample t-test; **Fig. 1D**). This is likely related to the smaller amplitude of gBOLD activity itself (*p* = 5.0*×*10^-4^, two-sample t-test; **Fig. 1E**). These two gBOLD metrics in the intermediate menopausal group fall between those of the other two groups, resulting in their significant linear relationships with the menopause age (gBOLD-CSF coupling: *p* = 0.039; gBOLD amplitude: *p* = 0.0015; ordinal regression). These observed gBOLD differences between the earlier and later menopausal groups cannot be attributed to head motion (**Fig. S2**).

**Table 1.** The demographics of Menopause groups.

| $N = 90$ | Earlier Menopause | Intermediate Menopause | Later Menopause | ANOVA p-value |
| --- | --- | --- | --- | --- |
| Number of subjects | 30 | 30 | 30 |  |
| Age at MRI (SD) | 58.1 (3.7) | 58.2 (4.0) | 60.0 (3.7) | 0.11 |
| Age at menopause (SD) | 45.5 (3.5) | 50.4 (0.6) | 54.0 (1.4) | $6.5 \times 10^{-25}$ |
| Mean education years (SD) | 14.9 (2.0) | 15.6 (1.5) | 15.8 (1.5) | 0.15 |
| Mean BMI (SD) | 28.3 (6.4) | 25.9 (4.4) | 26.8 (4.1) | 0.18 |

**Figure 1.**
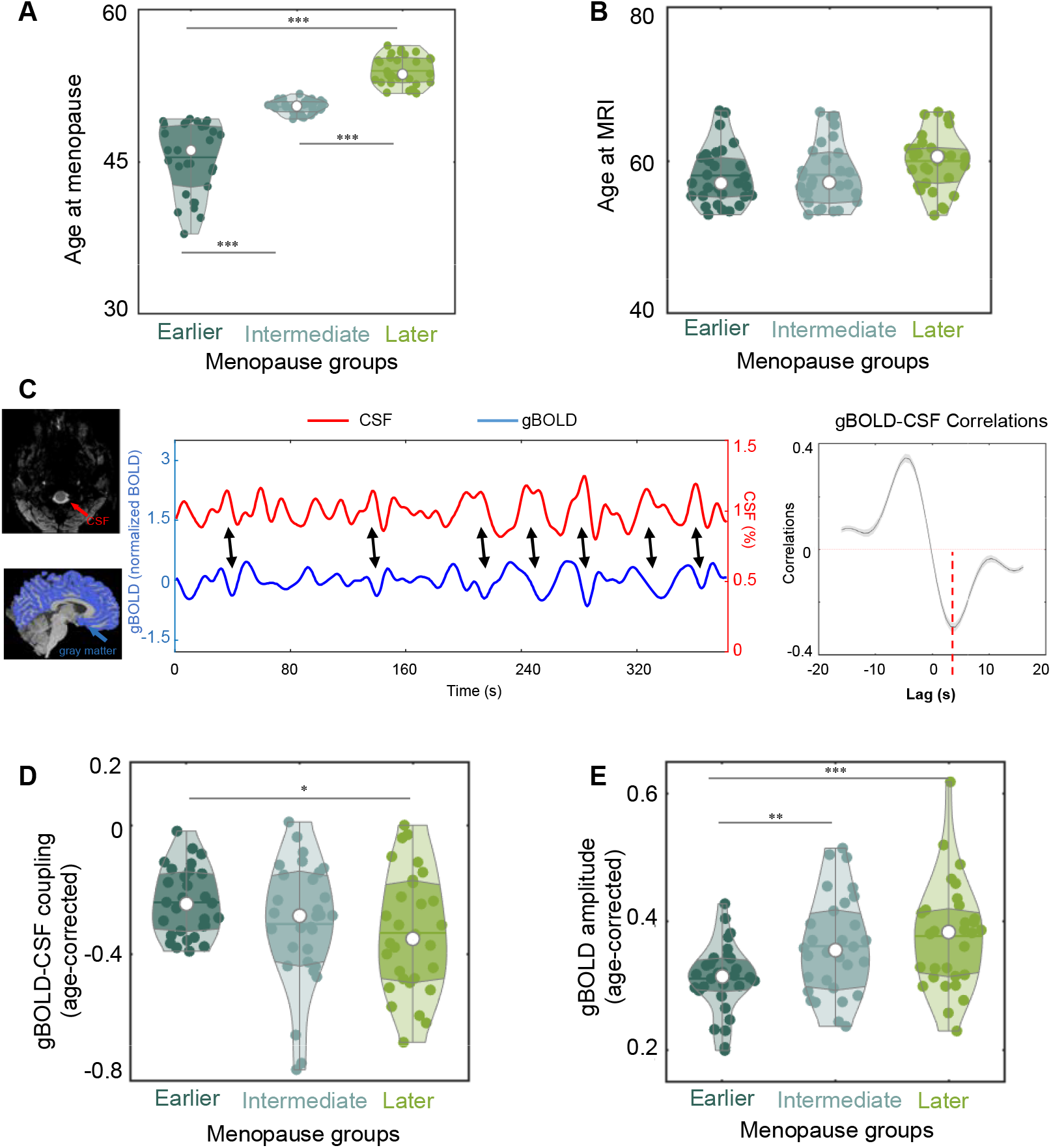
gBOLD-CSF coupling and gBOLD amplitude across women groups with different menopausal ages. (**A**) Three equally sized groups (N = 30 for each) based on age at menopause (earlier vs. intermediate: p=1.9×10^-10^, earlier vs. later: p=6.2×10^-18^, intermediate vs. later: p=9.5×10^-18^, two-sample t-test). (**B**) The three groups exhibit no significant differences (p = 0.11, ANOVA) in their age at MRI. (**C**) The gBOLD and CSF signals from one representative subject show corresponding changes (indicated by black arrows), resulting in significant correlations as seen in their mean cross-correlation function across 90 subjects (right panel). Individual subjects’ coupling strength is estimated as the correlation at the +3.2-seconds lag (red dashed line), corresponding to the negative peak of the mean cross-correlation function. Shaded regions indicate the range within one standard error of the mean. (**D**) The earlier menopausal group exhibits significantly weaker gBOLD-CSF coupling (age-corrected) than the later menopausal group (p=0.017, two-sample t-test). (**E**) The women with earlier menopause also show the smallest gBOLD amplitude (age-corrected) among all three groups. (earlier vs. later: p=5.0×10^-4^, two-sample t-test; earlier vs. intermediate menopause, p=0.0044, two-sample t-test). The significant changes are marked by asterisks (^*^: 0.01<p<0.05; ^**^: 0.001<p<0.01, ^***^: p<0.001).

### gBOLD differences in earlier menopausal women are mostly found in higher-order brain regions

We then examined the topology of gBOLD activity, which has been linked to the spreading pattern of Aβ in the early stages of AD^28^. We estimated the contribution of regional BOLD signals (rBOLD) to gBOLD amplitude through gBOLD presence (i.e., rBOLD-gBOLD correlations) and to gBOLD-CSF coupling via rBOLD-CSF couplings (see Methods for detail). The gBOLD presence maps showed significant differences between the earlier and later menopausal groups, predominantly in the frontal regions, cingulate cortex, and parietal regions (**Fig. 2A**). These regions overlap with the DMN and frontoparietal network (FPN)^35^ (**Fig. 2C**), which are associated with high principal gradient (PG) scores representing high hierarchical levels^36^. The finding suggests these higher-order regions are more de-synchronized from gBOLD activity in the earlier menopausal group compared to the later menopausal group. The comparison of rBOLD-CSF coupling between the two groups revealed a similar pattern, showing that a similar set of higher-order brain regions show weaker coupling to CSF flow in the earlier menopausal group, compared to the later menopausal group (**Fig. 2B**).

**Figure 2.**
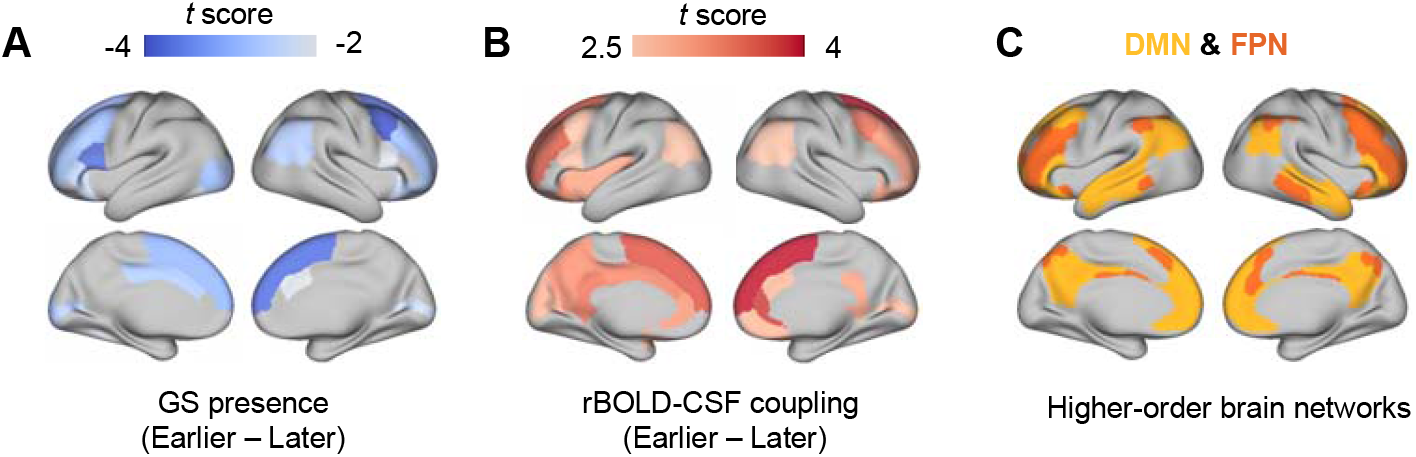
Comparison of gBOLD presence and rBOLD-CSF coupling between the earlier and later menopausal groups. The gBOLD presence (**A**) and rBOLD-CSF coupling (**B**) show significant differences between the later menopause and the earlier menopause, primarily in the higher-order brain regions belonging to the default mode network (DMN) and frontoparietal network (FPN) (**C**)^35^. Only brain parcels showing a significant difference (p < 0.05, two-sample t-test) are color-coded according to their t-scores in (**A**) and (**B**).

### Changes in gBOLD activity as propagating waves in earlier menopausal women

The gBOLD peaks are often manifested as propagating waves across cortical hierarchies^23,24^. We then examined whether gBOLD waves, which are expected to contribute significantly to the gBOLD and its coupling with CSF inflow, are modulated in earlier menopausal women. We identified gBOLD propagating waves^23,28^ that appear as tilted bands in the time-position graph along the PG direction and are more intuitive on the brain surface (**Fig. 3A-B** and **Fig. S3**). In earlier menopausal women, SM-to-DMN waves showed significantly weaker activations in higher-order brain regions compared to the later menopausal group (**Fig. 3C**). But this difference was largely absent in the DMN-to-SM propagating waves or in gBOLD peaks without propagating activity (**Fig. S3A** and **S3B**).

**Figure 3.**
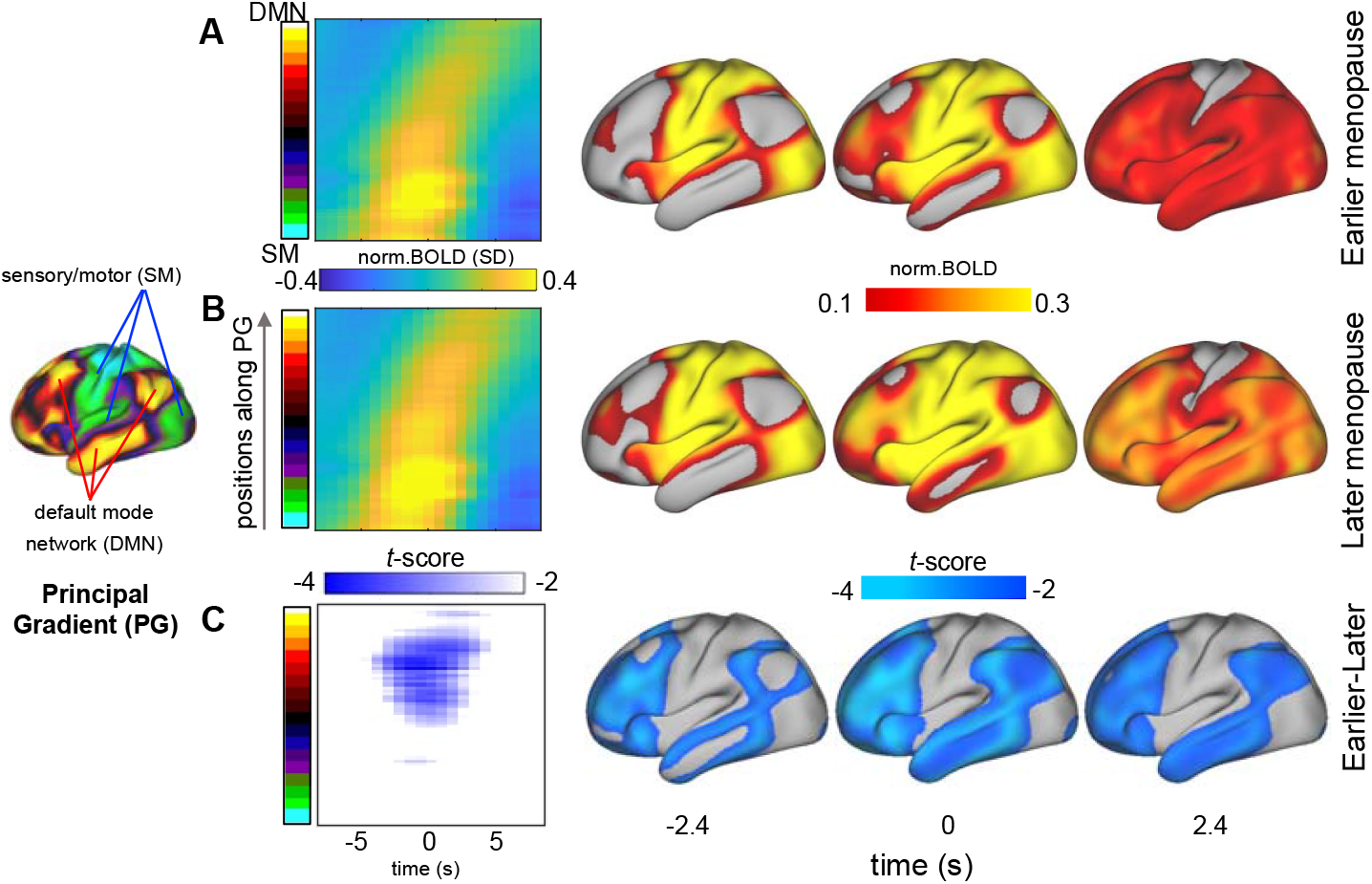
SM-to-DMN propagating waves disengage from higher-order regions in earlier menopausal women. The gBOLD waves propagating from the sensory-motor (SM) areas to the default mode network (DMN) are averaged for the earlier (**A**) and later (**B**) menopause groups, with their differences shown in (**C**). The average SM-DMN waves are manifested as tilted bands in the time-position graph (2nd column) and more visually intuitive when mapped onto the brain’s surface (3^rd^ to 5^th^ columns). A two-sample t-test was used to compare the propagating waves from the two groups in the (**A**) and (**B**).

### Memory performance correlates with gBOLD wave activations in higher-order regions

We then investigated how the observed gBOLD changes relate to memory function. The HCP-A dataset includes performance scores from the Picture Sequence Memory Test (PSMT) and the Rey Auditory Verbal Learning Test (RAVLT), which primarily assess memory functions, particularly episodic memory. The earlier menopausal group exhibited significantly lower PSMT scores than the later menopausal group (*p* = 0.008, two-sample t-test) (**Fig. 4A**), consistent with previous observations^37^. Among the earlier menopausal women, those with higher PSMT scores tended to have more SM-to-DMN waves during sessions of the same duration (*r* = 0.46, *p* = 0.02) (**Fig. 4B**). The same analysis on RAVLT scores found a similar, but marginally significant (*r* = 0.34, *p* =0.083) correlation with the number of SM-to-DMN waves (**Fig. S4A-B**). In comparison, the two memory scores are not correlated with the number of DMN-to-SM waves (*r* = 0.064, *p* = 0.76, for the PSMT; *r* = 0.036, *p* = 0.86, for the RAVLT) (**Fig.S4 C-D**), or with the number of SM-to-DMN waves among later menopausal women (*r* = −0.080, *p* = 0.68, for the PSMT; *r* = 0.017, *p* = 0.93, for the RAVLT) (**Fig. S4E** and **S4G**). Lastly, a direct correlation between the activation level of SM-to-DMN waves and PSMT score in earlier menopausal women revealed significant (*p < 0*.*05*) associations in higher-order regions at the late wave phase (**Fig. 4C**). Subjects with lower PSMT scores tended to have weaker activations in these regions at the SM-to-DMN waves (**Fig. 4C**, middle). Very similar results were also obtained for the RAVLT scores (**Fig. 4C**, right). In contrast, the associations between memory functions and wave dynamics are absent among the later menopausal women (**Fig. S5**).

**Figure 4.**
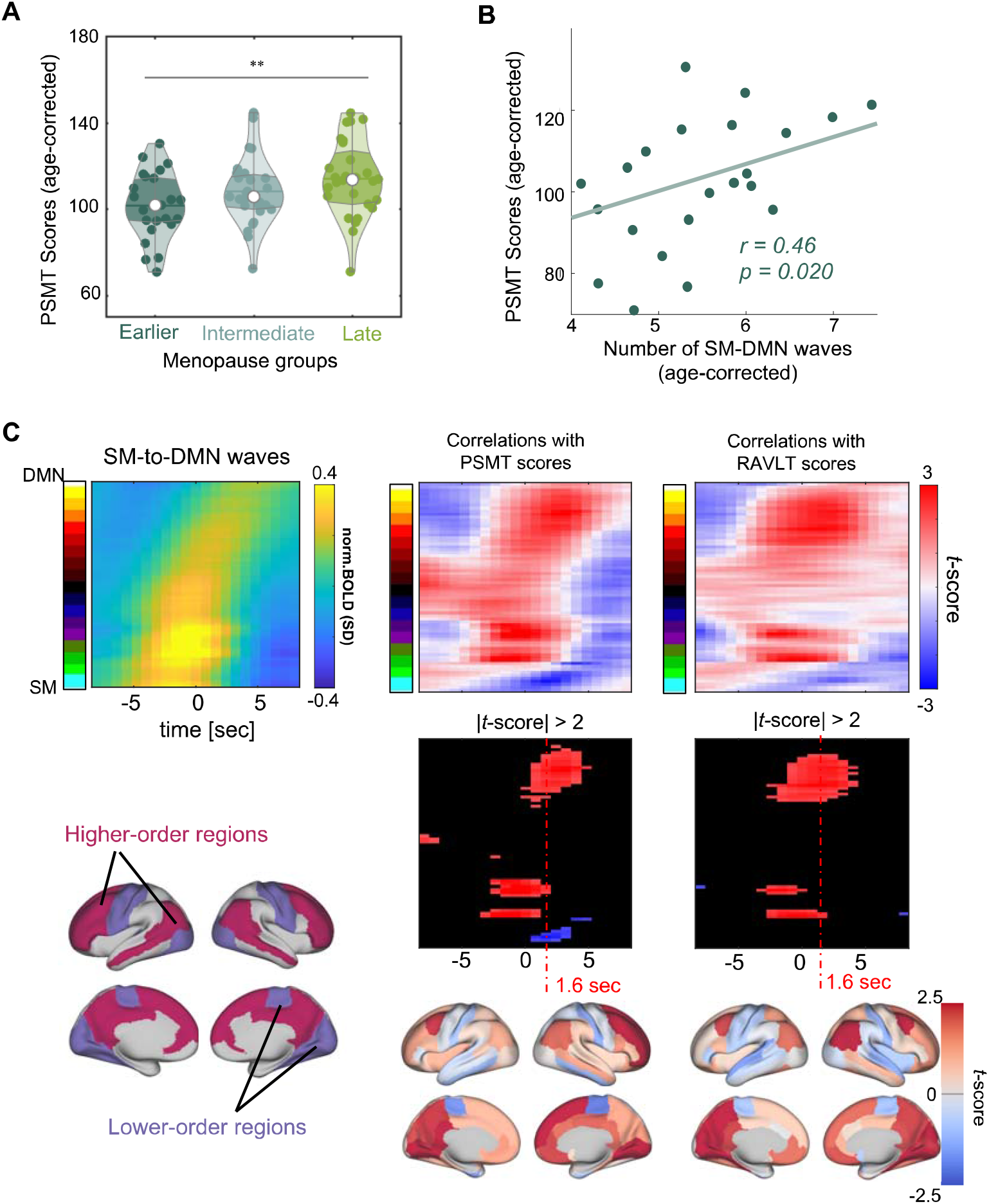
Weaker gBOLD wave activations in higher-order brain regions are correlated with poorer memory performance in earlier menopausal women. (**A**) The earlier menopausal group (N = 25) demonstrates significantly poorer performance on the Picture Sequence Memory Test (PSMT) than the later menopausal group (N = 28) (p = 0.008, two-sample t-test). The PSMT scores have been adjusted for age at MRI. (**B**) The number of SM-DMN propagation is significantly correlated with PSMT scores in the earlier menopausal group (r = 0.46, p = 0.020). (**C**) The SM-to-DMN propagating waves are averaged in the earlier menopausal group (top left). Across earlier menopausal women, SM-to-DMN waves are significantly (p < 0.05) correlated with PSMT scores in higher-order brain regions during the late wave phase (middle column). Their correlations (converted to t-score) at the +1.6-second wave phase are mapped onto the brain surface (bottom middle) and show network patterns corresponding to the higher- and lower-order brain regions defined previously (bottom left) ^28,35^. Similar results were also observed for RAVLT scores (right column).

## Discussion

The present investigation, using resting-state fMRI data from the Human Connectome Project - Aging, finds that women who experienced earlier menopause (in their 30s and 40s) exhibit weaker resting-state infra-slow global brain activity in later life compared to women who experienced menopause in their 50s. The weaker global activity is reflected by the smaller amplitude of gBOLD signal and its weaker coupling with CSF flow, which have been previously linked to AD pathologies, particularly Aβ accumulation^28,30^. Importantly, earlier menopausal women showing weaker gBOLD activation at higher-order regions performed worse on two episodic memory tests. These novel findings highlight brain activity changes linked to early menopause, suggesting a potential mechanism underlying memory decline and the increased risk of dementia in early menopausal women.

### The association between gBOLD activity and episodic memory in earlier menopause women

The observed correlation between gBOLD activity and memory performance in earlier menopause women is consistent with the known functional relevance of the infra-slow global activity. The cross-hierarchy propagation of this global activity resembles the cross-layer information flow required for optimizing large-scale artificial neuronal network models^38,39^, suggesting its potential role in learning and memory. The gBOLD waves also critically involve the neuromodulatory systems (particularly the cholinergic and noradrenergic systems) of memory relevance ^40^, as evident by strong and specific deactivations in their key hubs, including the nucleus basalis (NB) of the basal forebrain and the locus coeruleus (LC) of the brainstem^23,41^. During the “offline” resting state, the spiking cascades, the manifestation of this global brain activity at the single neuron level in mice, are indeed accompanied by changes in the rate of hippocampal sharp wave ripples (SPW-Rs) and replay events, which are important for memory retrieval and consolidation^21,22^. During “online” task performance, the ripple rate (in mice) and memory recalling (in human) are systematically modulated across the cycle of these global dynamics, with the DMN-activated phase being associated with enhanced memory retrieval^20,27^. Thus, changes in gBOLD activity, particularly those at the DMN-activated phase, have potential to influence both “offline” memory consolidation and “online” memory retrieval and thus are related to episodic memory performance in earlier menopausal women as estimated by the PSMT and RAVLT^42,43^.

The gBOLD-memory correlation was observed only in earlier menopausal women, but not in the other groups. This may be because the earlier menopausal group we defined has a much wider range of menopausal age (38.0–49.2 years) than other studies, and this range includes both normal (45–49 years) and early (<45 years) menopause as traditionally defined^44^. Weaker gBOLD-CSF coupling has been associated with lower cognitive scores in patients with mild cognitive impairment (MCI), AD, and Parkinson’s Disease (PD)^28–30^. These previously observed coupling-cognition associations may actually be attributed to changes in gBOLD activity/waves.

### Potential vascular contributions

The above discussion on the neural aspect of gBOLD activity does not preclude potential vascular contributions. The gBOLD activity is associated with strong physiological and vascular modulations, even at peripherals^45,46^. In addition to local neurovascular coupling, a global mechanism involving the sympathetic system may contribute to gBOLD signals, particularly its negative peak^45^. The subcortical nuclei, including the NB and LC that showed significant deactivations at gBOLD waves, could be the key structures mediating vascular effects associated with changes in gBOLD activity^23,41,47^. Thus, the gBOLD activity reduction and associated memory decline in earlier menopause women may also be associated with vascular dysfunctions, including those observed previously^48^. For example, as women traverse menopause, there is accelerated stiffening of the large elastic (aortic and carotid) arteries and increases in abdominal and thoracic perivascular fat^49^. Each of these vascular risk factors has been linked to faster cognitive decline and a higher risk of dementia in later life^50^. Future studies on the link between the infra-slow brain activity and vascular factors would likely enhance our understanding of why women who experience early menopause become more susceptible to cognitive decline later in life, and lead to targeted countermeasures.

### gBOLD-CSF coupling alterations in earlier menopause: harbinger of early Aβ pathology?

The gBOLD changes in earlier menopausal women, particularly its preferential decline in higher-order brain regions, resemble those observed in the early stages of AD^51^, which have been associated with the accumulation and spreading of Aβ^28^. In addition to arousal indices and hippocampal SWRs, global brain activity is coupled to CSF flow^32,46,52^, suggesting its possible role in perivascular clearance. This coupling is particularly strong during sleep^32^, further aligning with the characteristics of a recently proposed glymphatic system for brain waste clearance^33,53,54^. The cholinergic and noradrenergic systems may alternatively dilate and constrict arteries across the cycle of the global brain activity, as evident by the NB and LC deactivation separated by a half cycle, to generate vasomotion that drives the CSF flow needed by the glymphatic system^47^. Subsequent studies indeed found significant correlations between the gBOLD-CSF coupling and various AD pathologies, including Aβ and tau accumulation^30,55^. The gBOLD-CSF coupling changes in the early stage of Aβ pathology were later attributed to changes in gBOLD activity itself^28^. Most intriguingly, the diminished engagement of gBOLD activity in higher-order DMN areas, which is at least partly due to SM-to-DMN waves failing to reach these regions, was found to account for the preferential Aβ deposition in these brain areas^28^. These topologically biased changes resemble what we observed in earlier menopausal women in the present study. Early menopausal women have been found to face an increased risk of Aβ accumulation^37^ and AD^14^, and tend to have greater tau accumulation at the same level of Aβ^56^. Changes in gBOLD activity and its coupling to CSF dynamics might reflect impaired waste clearance, which could contribute to the increased risk of toxic protein accumulation and thus dementia in this population.

### Possible mediators of altered global brain activity in earlier menopausal women

Estrogen may be the key linking these gBOLD changes to existing hypotheses and theories about early menopause. The menopausal transition is marked by a sweeping decline in the productive of ovarian hormones – up to 90% for both estrogen and progesterone^9^, and women who experienced menopause earlier appear to have lower lifetime exposure to estrogen than those who underwent it later^57^. Additionally, postmenopausal women exhibit higher rates of Aβ deposition compared with premenopausal women and men after adjusting for age^58^. Reduced estrogen levels have been the primary hypothesis for the increased risk of dementia in early menopausal women, presumably due to its neuroprotective effects, such as inhibiting Aβ plaque formation, reducing oxidative cell damage^59^, and vascular protection^60^. Importantly, estrogen also enhances the cholinergic system by increasing acetylcholine levels, modulating cholinergic receptors, and protecting cholinergic neurons^40,61–64^. Interestingly, the cholinergic system plays a crucial role in the infra-slow global brain activity. The basal forebrain NB, the main subcortical cholinergic center, exhibits specific and strong deactivations during gBOLD SM-to-DMN waves^23,41^. Pharmaceutical deactivation of the NB in one brain hemisphere effectively suppressed gBOLD activity on the ipsilateral side, confirming its causal role in gBOLD regulation^65^. Thus, estrogen appears capable of influencing gBOLD activity via cholinergic pathways. It is also worth noting that DMN hubs, where we observed the major gBOLD changes in the ealier menopausal group, also coincide with brain regions characterized by high estrogen receptor density^66–69^.

Estrogen changes associated with menopause have been linked to resting-state fMRI connectivity previously. Compared to premenopausal women, postmenopausal women with lower estrogen showed weaker functional connectivity between the left medial orbitofrontal cortex and the right thalamus^70^. Lower estrogen levels in postmenopausal women have also been associated with weaker DMN-related functional connectivity^71^. In perimenopausal women, regional homogeneity in the superior frontal gyrus has also shown a positive correlation with estrogen levels^72^. The gBOLD activity is expected to significantly influence resting-state connectivity in a complicated way^73–75^. The Aβ-associated decrease in DMN connectivity in AD has been shown to reflect DMN-dominant gBOLD changes^28^. Thus, it is possible that estrogen-associated–changes in functional connectivity, particularly those in the higher-order DMN regions, may originate from the DMN-dominant gBOLD changes we observed here in the earlier menopausal women. To clarify this relationship, future studies should investigate this possibility by examining the relationship between more lifetime estrogen exposure and gBOLD activity.

### Study limitations

One limitation of this study is that there was no control over the availability or completeness of the HCP-A dataset, particularly regarding key variables such as hormone therapy (HT) use and history, lifestyle behaviors (habitual physical activity level, dietary practices, etc.), detailed medical history to understand potential drivers of early menopause (e.g., cancer, reproductive surgeries), prior brain trauma, etc. Hormone therapy has been shown to lower the risk of Alzheimer’s disease^76^ and increase cholinergic activity^77^, potentially influencing gBOLD activity. Although both natural menopause and surgical menopause are associated with an increased risk of dementia^37,78^, the risk linked to surgical menopause tends to be more severe^78^. Therefore, future research should explore the effects of hormone therapy on gBOLD activity and its interaction with the age of menopause.

## Conclusions

In summary, our study found that earlier menopausal-onset women exhibit weak global brain activity measured by multiple gBOLD metrics, which have been previously associated with cognitive decline and Aβ accumulation in the early stages of AD^28,30^. The gBOLD change is mainly characterized by reduced activation in higher-order DMN/FPN regions, which correlates with poorer episodic memory performance, a factor thought to play a key role in cognitive aging^18^. These findings suggest a possible mechanism underlying the increased risks of AD and dementias in early menopausal women.

## Acknowledgments

This work was supported by the Brain Initiative award (1RF1MH123247-01) and the NIH R01 award (1R01NS113889-01A1).

## Data and materials availability

Data and code used to generate the main results are available upon request.

## Materials and Methods

### Participants and study data

This study used fMRI and behavioral data from the HCP-Aging project (https://www.humanconnectome.org/study/hcp-lifespan-aging)^34^. All participants provided written informed consent, and investigators at each HCP-A participating site obtained ethical approval from the corresponding institutional review board (IRB). The use of de-identified data from the HCP-A and the sharing of analysis results have been reviewed and approved by the Pennsylvania State University IRB (IRB#: STUDY00008766) and also strictly followed the National Institute of Mental Health (NIMH) Data Archive-data use certification (DUC).

We identified 124 healthy postmenopausal females (36.5∼78.1 years old) with available information on their age at menopause, who also completed all four resting-state fMRI sessions. Here, menopause status is determined by the question of “whether having no period for 12 months” in the “menstrual cycle” data (mchq01.txt). The age at menopause was determined by the participant’s age at MRI and the answer to “When was your last period?”. Participants with abnormal responses (e.g., reporting a last period later than their age at MRI) and missing values were excluded from analyses.

To group the 124 subjects by menopausal age while minimizing confounding with age at MRI, we first divided them into two groups according to the U.S. average menopause age of 51^79^ (**Fig. S1**), and found the overlapped range of their age at MRI. We then excluded 34 subjects whose age at MRI fell outside this range (52.9 to 66.5). The remaining 90 subjects were subsequently divided into three equally sized groups—earlier, intermediate, and later menopause—based on their age at menopause (**Fig. 1A**). The three groups showed no difference in their age at MRI (*p*=0.11, ANOVA; **Fig. 1B**).

### Behavioral Data

We used the measurements of episodic memory function provided by the HCP-A project. The HCP-A study measured episodic memory using the NIH Toolbox Picture Sequence Memory Test (PSMT) and the Rey Auditory Verbal Learning Test (RAVLT)^80^. The PSMT evaluates episodic memory by requiring participants to recall sequences of illustrated objects and activities. Participants were instructed to recall the sequence of images shown during two learning trials, with sequence length varying by age from 6 to 18 pictures. Performance was determined by the number of correctly recalled adjacent picture pairs, with the maximum score equal to one less than the total number of items in the sequence. This study used the “unadjusted scaled score for PicSeq subtest” variable as the final PSMT score. The RAVLT assesses verbal learning and memory, including components of short-term memory. The test consists of multiple trials involving two word lists. In the first five trials, subjects hear and then recall a list of unrelated words (List A). Novel words not on List A and repetitions are also tracked and scored. In the sixth trial, participants are presented with an interference list (List B) and asked to recall it. During the final trial, participants recall words from List A without hearing them again. Recall of interfering words—List A during trial 6 and List B during trial 7—is also monitored and scored. We used the “RAVLT Short Delay Total Correct” variable to quantify RAVLT task performance. Subjects with missing or abnormal task performance scores (coded as 999 or NaN) were excluded from the analysis. Consequently, for the PSMT task, the final sample included 25 earlier menopausal, 27 intermediate menopausal, and 28 later menopausal individuals. For the RAVLT task, the respective group sizes were 27, 30, and 29 participants.

### Image acquisition and preprocessing

Resting-state fMRI data were collected using 3T MR scanners (Siemens Medical Solutions, Siemens, Erlangen, Germany) following a standardized protocol^34^ across four acquisition sites: Washington University St. Louis, University of Minnesota, Massachusetts General Hospital, and University of California, Los Angeles. Data analysis was conducted by researchers at Oxford University. Each subject underwent two sessions of eyes-open resting-state fMRI (REST1 and REST2) over one or two days, depending on the site. Each session consisted of two 6.5-minute runs with opposite phase-encoding (PE) directions: REST1 Anterior-Posterior, REST1 Posterior-Anterior, REST2 Anterior-Posterior, and REST2 Posterior-Anterior. Following the fMRI scans, a T1-weighted structural MRI session was conducted using the MPRAGE sequence with the following parameters: echo time (TE) = 1.8/3.6/5.4/7.2□ms (multi-echo), repetition time (TR) = 2,500□ms, field of view (FOV) = 256□×□256□mm^2^, matrix = 320 × 300, number of slices = 208, voxel size = 0.8□×□0.8□×□0.8□mm^3^, and flip angle = 8° (Harms et al., 2018). The T1-weighted MRI provided whole brain and CSF volume data and was used for anatomical segmentation and registration. For the rsfMRI acquisition, 478 fMRI volumes were collected using a multiband gradient-recalled echo-planar imaging (EPI) sequence with the following parameters: TR/TE = 800/37 ms, flip angle = 52°, field of view (FOV) = 208 mm, matrix = 104 × 90, 72 oblique axial slices, 2 mm isotropic voxels, and a multiband acceleration factor of 8.

For the gBOLD and CSF signals, we utilized data from the previous study^31^. RsfMRI data were preprocessed for gBOLD and CSF signal extraction using FSL (version 5.0.9; https://fsl.fmrib.ox.ac.uk/fsl/fslwiki/FSL)^81^ and AFNI (version 16.3.05; https://afni.nimh.nih.gov/)^82^, following the protocol outlined in previous study^30^. The general fMRI preprocessing procedures included motion correction, skull stripping, spatial smoothing (full width at half maximum (FWHM) = 4mm), temporal filtering (bandpass, approximately 0.01 to 0.1 Hz), and the co-registration of each fMRI volume to corresponding T1-weighted structural MRI^29,30^. In line with previous work^28–30^, motion parameters were not regressed out to avoid attenuating the gBOLD signal, as recent findings suggest that head motion and its “apparent” effects on rsfMRI signals^83^ may actually be by-products of gBOLD activity^23,84^. Structural image preprocessing, including spatial normalization and skull stripping, was performed using FSL.

To quantify propagating waves and gBOLD presence, we used the pre-processed surface-based fMRI released by HCP-Aging. Specifically, the rsfMRI data were processed based on an established pipeline^85^ using FSL^86^, FreeSurfer^87^, and HCP workbench^88^. Additionally, the HCP FIX-ICA denoising pipeline was applied to remove artifacts. To improve intersubject registration, the multimodal surface matching registration was used in the HCP dataset^89,90^. The rsfMRI cortical surface data were mapped to the standard HCPfs_LR32k surface mesh, with each hemisphere containing 32,492 nodes (totaling 59,412 nodes excluding the non-cortical medial wall). We further temporally filtered the data using a bandpass filter between 0.01 and 0.1 Hz, and standardized the signal at each vertex by subtracting the mean and dividing by the standard deviation.

### Extraction and computation of gBOLD and CSF signals

The gBOLD and CSF signals were extracted following previously established methods^29,30^. Gray matter regions were defined using the Harvard-Oxford cortical and subcortical structural atlases (https://neurovault.org/collections/262/). To avoid spatial blurring from the MNI-152 space^32^, we registered the gray matter mask back to the native space of individual subjects. We then spatially averaged the Z-normalized fMRI signals across all gray matter voxels to obtain the gBOLD signal. The CSF inflow signal was extracted at the bottom slice of fMRI acquisition from a region positioned below the cerebellum, which is consistent across all subjects, to maximize sensitivity to CSF inflow effects^32,91,92^. The CSF region of interest (ROI) was defined from the preprocessed fMRI data in each participant’s native space. ROIs were manually drawn based on T2^*^-weighted fMRI, where CSF appeared brighter than surrounding tissues, and their locations were confirmed using higher-resolution T1-weighted MRI images.

To quantify gBOLD-CSF coupling, the cross-correlation function was first computed between the extracted gBOLD and CSF signals for each fMRI session of every participant. This function assessed Pearson’s correlation at time lags ranging from −16 to +16 seconds. For each participant, the cross-correlation functions from the four individual fMRI runs (AP1, PA1, AP2, and PA2) were averaged. These cross-correlation functions were then averaged across all 90 subjects (**Fig. 1C**). The negative peak at the lag of +3.2 seconds in **Fig. 1C** reflects the peaked coupling relationship between the gBOLD and CSF signal. Following previous studies^29,30^, the gBOLD-CSF coupling for each participant was determined by her/his cross-correlation function at a time lag of +3.2 seconds.

### Quantification of gBOLD presence and rBOLD-CSF coupling

To quantify gBOLD presence and rBOLD-CSF coupling, we first extracted regional BOLD signals from the surface-based fMRI using the DKT-68 parcellation^93^. The fMRI BOLD signals were averaged within each parcel to obtain the regional BOLD signal for each cortical region. Following the approach outlined in a previous study^94^, the regional BOLD signal of each region was then correlated (Pearson’s correlation) with the mean BOLD signal across the entire cortical surface, resulting in gBOLD presence that represents the synchronization between global and regional brain activity. The regional BOLD signal of each region was also correlated with the CSF signal at the time lag of +3.2 seconds, and the resulting correlation quantifies the rBOLD-CSF coupling.

### Quantification of gBOLD propagation waves

We quantified gBOLD propagating waves following methods described in previous studies ^23,28^. Specifically, 59412 cortical surface vertices were first projected onto the direction of the principal gradient (PG), which was derived from a low-dimensional embedding of a mean connectivity matrix obtained from 820 subjects in a prior study^36^. The vertices were evenly divided into 70 positional bins (regions). Next, resting-state fMRI signals were averaged within each bin, resulting in 70 time series. The entire time course was segmented based on the troughs of the gBOLD signal (defined as the mean signal across all 59,412 cortical surface vertices). Within each time segment, the largest local peak was identified for each bin. Only time segments with a real “global” involvement (i.e., local peaks were detected in at least 80% positional bins) were used for further analyses. For each qualifying time segment, we computed the Pearson correlation between the relative timing of local peaks (time delay relative to the gBOLD peak) and their positions along the PG. Segments showing significant positive and negative correlations (p < 0.01) were classified as bottom-up (SM-to-DMN) and top-down (DMN-to-SM) propagating waves, respectively. Segments with no significant correlation (p>0.05) were classified as gBOLD peaks without propagation.

### Estimation and analysis of head motion

To examine whether head motion is related to the differences in gBOLD activity between the earlier and later menopausal groups, we first quantified head motion for each subject using the session-mean framewise displacement (mFD), following the approach outlined in previous studies^83,95^. Specifically, FD was calculated by summing the absolute values of all six translational and rotational realignment parameters at each time point as follows:

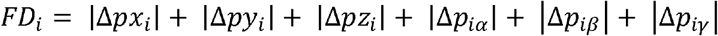

where [*px*_*i*_, *p* _*i*_, *p* _*i*_] are the three translational parameters and [*p*_*iα*_,*iβ*_*i*_, *p*_*iλ*_] are the three rotational parameters at the ith frame. |Δ*px*_*i*_| *px*_*(i+l)*_ - *px*_*i*_| and the other parameters were calculated in the same way. mFD was compared across menopausal groups and correlated with gBOLD amplitude and. gBOLD-CSF coupling. In addition, the main results were reproduced with adjusting the mFD.

### Statistical analysis

Before evaluating the impact of menopausal timing on global brain activities and episodic memory task performance, we first regressed out the linear effect of age at MRI from these measures in the selected 90 subjects. Subsequently, we performed two-sample t-tests to compare these age-adjusted parameters between the earlier and later menopause groups. The same age correction procedure was also applied before analyzing the correlation between the number of SM-DMN propagations and episodic memory task performance in the earlier menopausal group. A linear regression model was then fitted to examine the association between episodic memory performance and propagation patterns, using age at MRI and task performance as predictor variables and propagation intensity as the response variable. A p-value smaller than 0.05 is considered statistically significant.

## Notes

### Competing Interest Statement

The authors have declared no competing interest.

### Summary of Updates

The author order has been updated. No other changes were made.

