## supplementary_figures for "Early menopause is associated with reduced global brain activity"

**Affiliations:**

431 Chemical and Biomedical Engineering Building

The Pennsylvania State University

University Park, PA 16802-4400

**Author Contributions:** Xufu Liu and Xiao Liu contributed to the conception, design of the work, and investigation; Xufu Liu, Yutong Mao, and Feng Han acquired and processed the data; Xufu Liu and Xiao Liu contributed to data analyses and visualization; Xiao Liu devoted the efforts to the supervision, project administration and funding acquisition; Xufu Liu, and Xiao Liu contributed to drafting the paper; Xufu Liu, Laura Pritschet, Liying Luo, David N. Proctor, and Xiao Liu contributed to editing and reviewing of the paper.

**Key Words:** Early menopause; Global brain activity; Resting-state fMRI; Infra-slow brain activity.

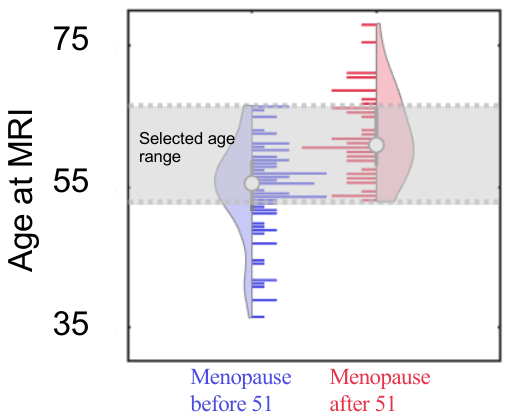

***Figure S1. Distributions of the age at MRI of all post-menopausal females with the information of menopausal age (n=124****). Blue: females who experienced menopause before 51 years old (n=77). Red: females who experienced menopause after 51 years old (n=47). To control the effect of age at MRI, we only focused on 90 females whose ages fell within the overlapping age range of the two groups (grey region, n=90).*

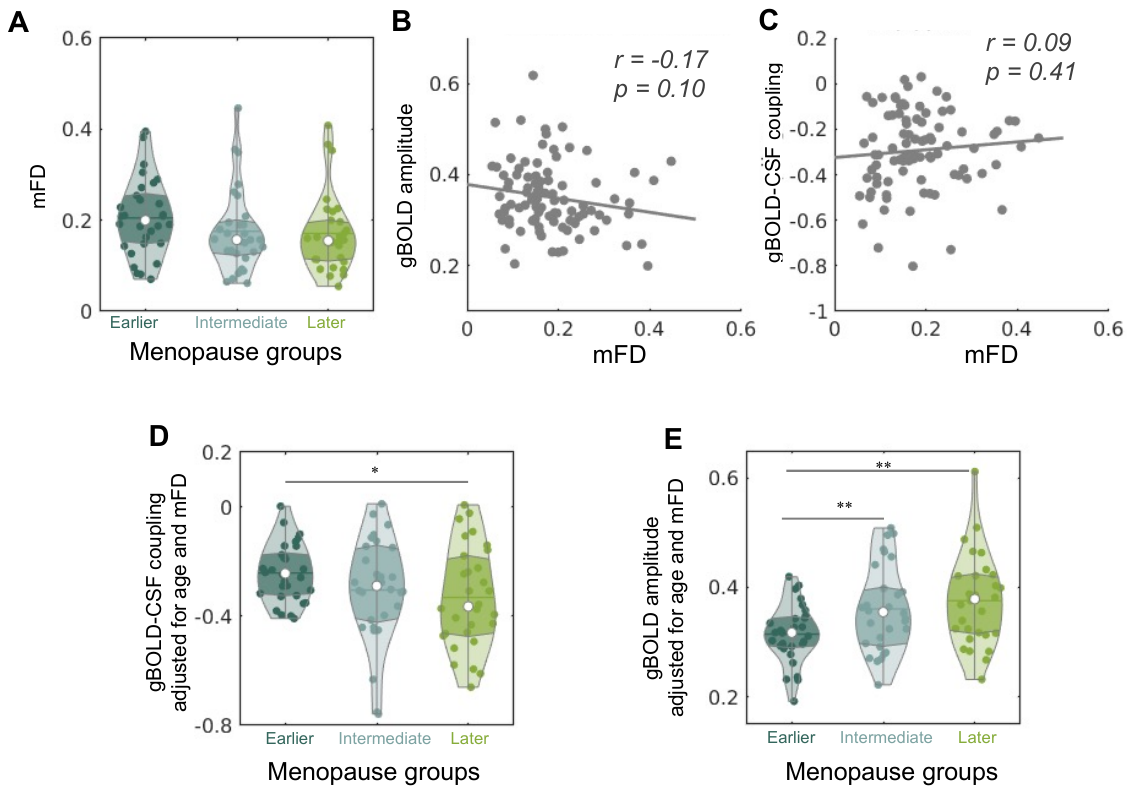

***Figure S2. Differences in gBOLD-CSF coupling and gBOLD amplitude between earlier menopausal and later menopausal groups, considering the effect of head motion.*** *The head motion is quantified by the mean framewise displacement (mFD).* *No significant differences in mFD were observed among the three menopausal groups (p = 0.26; ANOVA) (****A****). The mFD is not correlated with either gBOLD amplitude (****B****) or gBOLD-CSF coupling (****C****). After correcting for the mFD, the differences in gBOLD-CSF coupling (p=0.025, two-sample t-test) (****D****) and gBOLD amplitude (p=0.0011, two-sample t-test) (****E****) between the earlier and later menopausal groups, as well as the difference in gBOLD amplitude between the earlier and intermediate menopausal groups (p=0.0091, two-sample t-test), remain significant.*

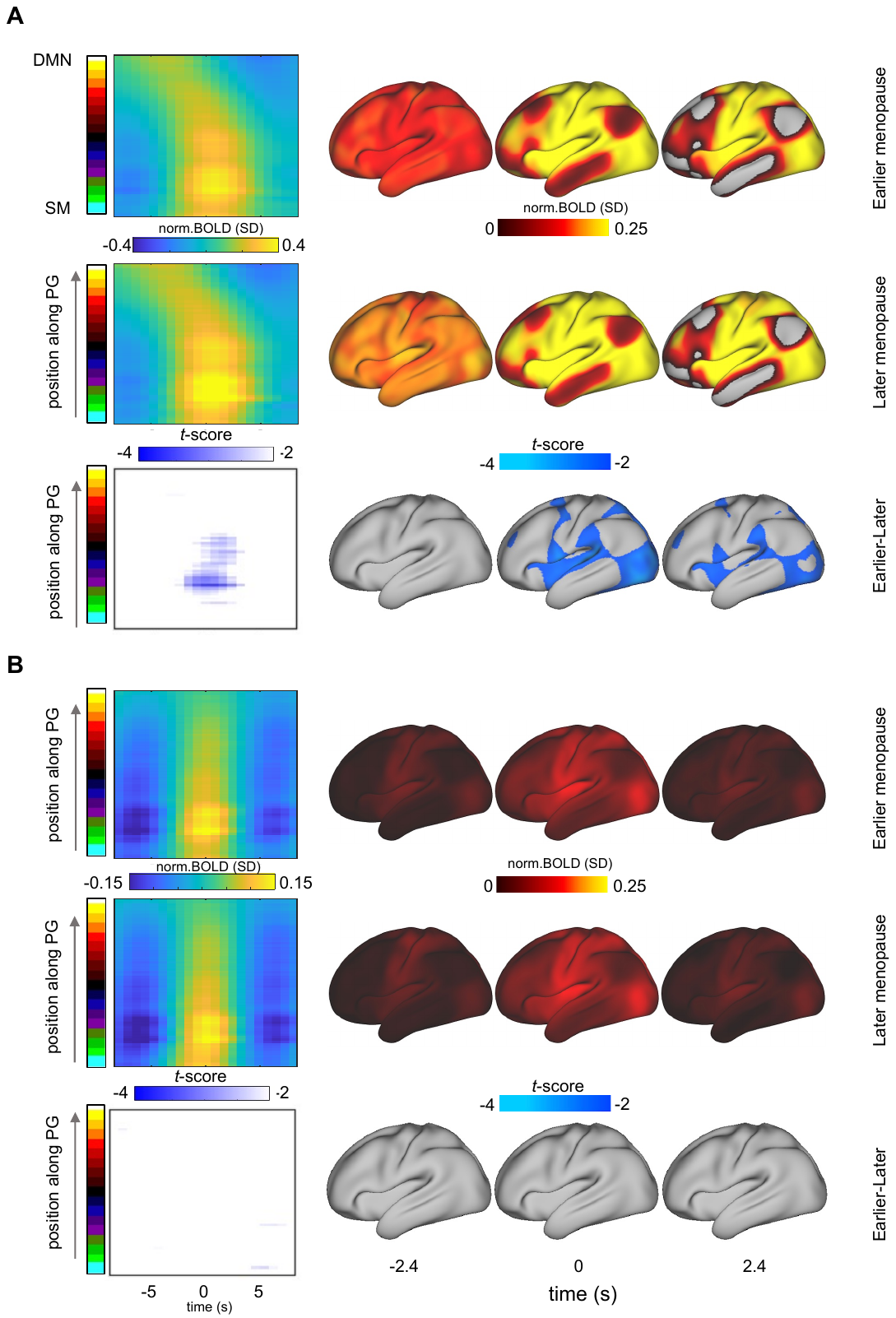

***Figure S3. DMN-to-SM propagating waves and gBOLD peaks without propagation do not exhibit significant differences between the earlier and later menopausal groups.*** *(****A****) The detected gBOLD waves propagating from the default mode network (DMN) regions to sensory-motor (SM) areas were averaged for the earlier (the first row) and later (the second row) menopausal groups, with their differences shown in the third row. (****B****) The gBOLD peaks without propagation were also averaged for earlier and later menopausal groups (the first two rows), with their differences shown in the third row.*

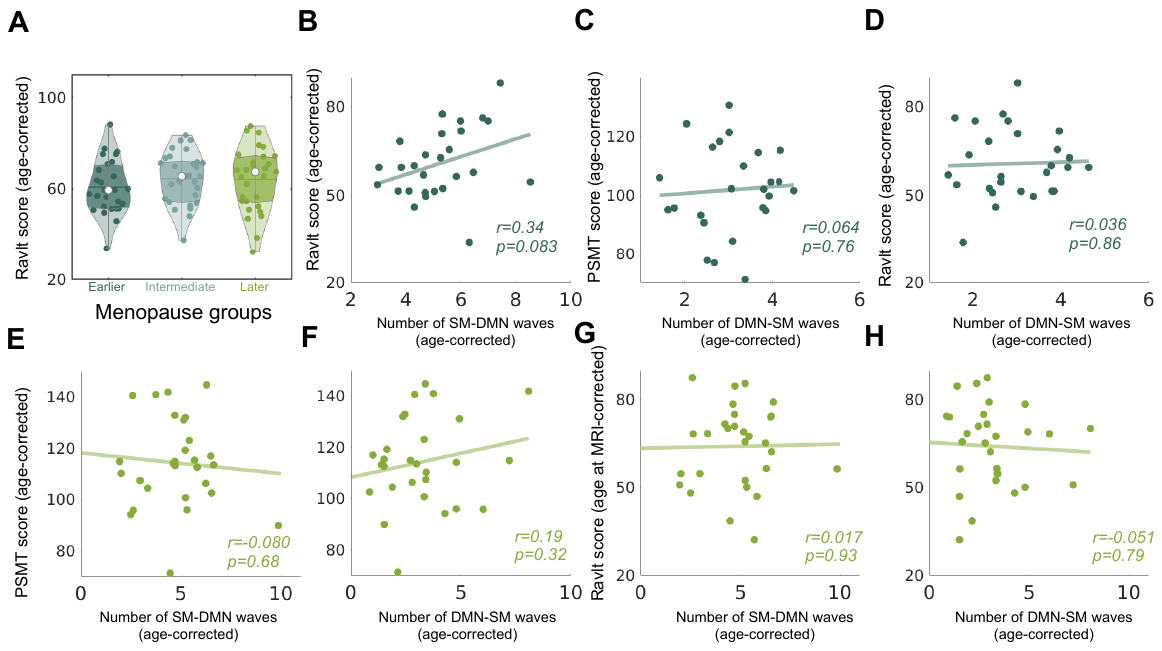

***Figure S4. Associations between the frequency of propagating waves and episodic memory task performance (PSMT and RAVLT).*** *(****A****)* *The mean RAVLT scores increased gradually from the earlier to intermediate and then to later menopausal groups, similar to the PSMT results, although no significant group differences were found (earlier vs. mid: p=0.23, earlier vs. late: p=0.34, intermediate vs. late: p=0.91, two-sample t-test). (****B****) The RAVLT scores showed a marginally significant correlation (r = 0.34, p = 0.08) with the number of SM–DMN propagating waves in the earlier menopausal group. No association was found between the number of DMN-SM waves with either RAVLT or PSMT scores in the earlier menopausal group (****C-D****), or between these memory scores and the number of propagating waves in the later menopausal group (****E-H****).*

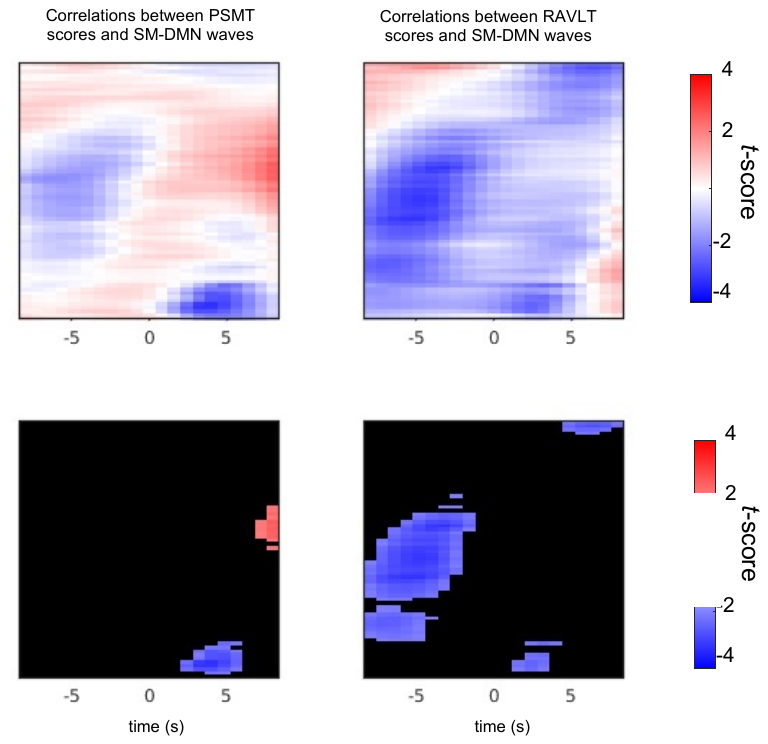

***Figure S5. Correlations between the SM-to-DMN waves and memory scores in the later menopausal group.*** *The* *t-score maps of the regression coefficients for PSMT (left) and RAVLT (right) scores on SM-to-DMN wave activations.*
